# Cell Cycle Delay, Pro-metaphase Arrest and C-metaphase Inducing Effects of Petroleum Ether Fraction of Leaf Aqueous Extract of *Clerodendrum viscosum* Vent

**DOI:** 10.1101/2020.11.29.402370

**Authors:** Sujit Roy, Lalit Mohan Kundu, Gobinda Chandra Roy, Manabendu Barman, Sanjib Ray

## Abstract

*Clerodendrum viscosum* is a traditionally used medicinal plant. The present study aimed to analyze cell cycle delay, pro-metaphase arrest, and c-metaphase inducing effects of the petroleum ether fraction (AQPEF) of leaf aqueous extract of *C. Viscosum* (LAECV). The LAECV was fractionated with petroleum ether and its metaphase arrest, cell cycle delay, and c-metaphase inducing activities were tested on *A. cepa* root tip cells. The AQPEF induced cell cycle delay, and colchicine like metaphase, c-metaphase, in *A. cepa* root tip cells. Thus, the present study explores AQPEF as an active fraction of LAECV having metaphase arresting, cell cycle delay, and c-metaphase inducing potentials.

---

*Clerodendrum viscosum* (Family: Lamiaceae) is widely distributed throughout Asia, Africa, Australia, and America. It is used in traditional, Ayurvedic, Homeopathic, and Unani medicine (Nandi and Lyndem 2015). The whole plant juice is used against worm infection, cough, itching, leprosy, scorpion sting, asthma, bronchitis, fever, *etc.* (Kirtikar and Basu 1991, Bhattacharjee 2011). The plant is well known for its effectiveness against rheumatism in Unani medicine (Singh *et al*. 1997). It is prescribed to treat postnatal complications, diarrhea, and fresh wounds in the Indian Homeopathic system (Hamilton 1997, Nadkarni and Nadkarni 2002). Its bark juice is used to relieve indigestion and abdominal pain. The different parts of this plant are used as a remedy for asthma, malaria, cataract, diseases of the skin, blood, and lung by the Indian Tribals of Chotanagpur plateau (Kirtikar and Basu 1991).

The plant’s ethanolic extracts have antioxidant, antimicrobial, hepatoprotective, wound healing, and antidiarrheal activities (Bhattacharjee *et al*. 2011). The anthelmintic activity was reported in leaf ethanolic, methanolic, and aqueous extracts against *Pheretima posthuma* (Islam *et al.* 2013). The leaves and roots have great potential against different microbial and fungal strains. Acetone and chloroform extracts of *C. viscosum* leaves have an inhibitory effect on the growth of *Shigella sp*., *Vibrio cholerae*, *Klebsiella pneumonia,* and *Pseudomonas aeruginosa, etc.* (Lobo *et al*. 2010) while ethanolic fraction has shown antifungal activity against *Aspergillus niger*, *A. flavus,* and *Candida albicans* (Modi *et al*. 2010). A saponin isolated from petroleum ether extract of leaves has analgesic activity (Sannigrahi *et al*. 2009). In vivo, antinociceptive activity of the methanolic extract is comparable to diclofenac sodium drug (Rahman *et al*. 1970). Reduction in CCl_4_ induced hepatotoxicity on rats after treatment with methanolic leaf extract reveals its hepatoprotective activity which is further supported by biochemical blood parameters (Sannigrahi *et al*. 2009). Successive ethanolic extract of *C. infortunatum* shows the highest antioxidant activity compared to petroleum ether and chloroform extracts, whereas significant wound healing activity was exhibited by petroleum ether and ethanol extract. These pharmacological effects were also correlated with the total phenolic content of the plant (Gouthamchandra *et al*. 2010). Leaf methanolic extract exerts a restoring effect on blood glucose levels after streptozotocin treatment (Arvind *et al*. 2002, Das *et al*. 2011). Allelochemicals from leaf aqueous extract have been found to harm the growth and germination of weeds in agro-ecosystem (Qasem and Foy 2001, Devi *et al.* 2013). A promising positive correlation is established between the plant’s parts and their insect repellent and insecticidal activity (Muh *et al.* 2014).

Acute toxicity test reveals that these plant parts are safe up to 2000 mg kg^−1^ body weight (Gupta *et al.* 2012). The crude leaf extracts contain phenolics viz. fumaric acids, acetoside, methyl esters of caffeic acids, terpenoids like clerodin, flavonoids such as apigenin, acacetin, scutellarein, quercetin, hispidulin, steroids such as clerodone, clerodolone, clerodol, clerosterol and some fix oils containing linolenic acid, oleic acids, stearic acid and lignoceric acid (Singhmura 2016).

In our previous study, we have reported that treatment with leaf aqueous extract of *C. viscosum* (LAECV) on root apical meristem cells of wheat and onion gave an increased metaphase frequency along with a reduction in mitotic index, antiproliferative, and apoptosis-inducing effects (Ray *et al*. 2012). The metaphase arrest and cell cycle delay-inducing effects were somewhat comparable to colchicine’s action (Ray *et al*. 2013, Kundu and Ray 2016, Roy and Ray 2017).

Colchicine inhibits spindle formation in cells which leads to generating signals delaying the transition from metaphase to anaphase (Salmon *et al*. 1984). Later on, when the concentration of colchicine decreases in the environment, the chromatids separate abnormally and the plant cells become polyploid (Caperta *et al*. 2006). Colchicine treatment to onion root apical meristem cells results in root growth inhibition, root’s swelling, haphazardly arranged condensed chromosomes, and increased frequency in metaphase (Ray *et al*. 2013). In another comparative study, colchicine and LAECV treatments on *A. cepa* root tip cells revealed similar cytotoxic effects (Kundu and Ray 2016). The metaphase arrest and cell cycle delay-inducing effects of LAECV raised the key question about its active principle(s) (Ray *et al*. 2013). Therefore, this study aims to fractionate the bioactive molecule(s) of LAECV with non-polar petroleum ether solvent and to test its antiproliferative and metaphase arresting activities. The LAECV was fractionated with petroleum ether and the extract fraction was tested for cell cycle delay, pro-metaphase arrest, and c-metaphase inducing effects on *A. cepa* root tip cells.

## Materials and methods

### Plant collection and extraction

The fresh *C. viscosum* leaves were collected from Burdwan University campus, West Bengal, India, and it was taxonomically identified by Professor A. Mukherjee, Department of Botany, The University of Burdwan. A voucher specimen (No. BUTBSR011) is maintained in the Department of Zoology, B.U., for future reference. Fresh leaves were washed in tap water, dried in shade, ground by Philips Mixer Grinder HL1605, and the obtained powder was stored in a tightly sealed container for further use. 100 g of this pulverized material was extracted in 2.5 L of boiling distilled water for 2-3 h and the extract was filtered with filter paper and the extract coded as leaf aqueous extract of *C. viscosum* (LAECV). The LAECV was fractioned by petroleum ether with the help of a magnetic stirrer for 10-12 h. After collection by siphoning, the solvent fraction (AQPEF) was concentrated by rotary vacuum evaporator and stored in a glass container.

### Culture and treatment for root growth retardation

1% sodium hypochlorite mediated surface-sterilized *A. cepa* bulbs were placed in 6-well plates containing distilled water and kept in the environmental test chamber for germination (25-27°C, humidity 50%). The 48h aged similar-sized *A. cepa* roots (2-3 cm root length) were treated with 12.5, 25, 50, 100, and 150 μg mL^−1^ concentrations of AQPEF in 1% DMSO continuously for 24, 48, and 72 h. The experiments were performed in triplicate.

### Cell cycle delay and c-metaphase inducing effects analysis

The 48h aged similar-sized *A. cepa* roots (2-3 cm) were treated with 50, 100, 150, 200 μg mL^−1^ of AQPEF and 150 μg mL^−1^ of colchicine for 2 and 4 h. The stock solution of the Petroleum ether fraction was prepared in DMSO for treatment. After 2 and 4 h exposure, 8-10 roots were fixed and processed for squash preparation following the standard procedure (Chaudhuri and Ray 2015). The remaining roots were allowed to grow further for another 16 h in distilled water (4h T+16h RS) and subsequently, root tips were fixed. The control group, which had not received any treatment, was maintained in distilled water simultaneously with the treatment groups. The treated and untreated root tips were fixed in aceto-methanol (3 parts methanol: 1part glacial acetic acid) for 24 h and then hydrolyzed for 10 min in 1 M HCl at 60□. The roots were stained with 2% aceto-orcein and finally squashed in 45% acetic acid (Sharma and Sharma 1999, Ray *et al.* 2013). The well-spread areas of squashed roots were focused under the bright field light microscope for observation and scoring the cell division phase and c-metaphase frequencies.

### Scoring and statistical analysis

In the case of squash preparation of *A. cepa* root apical meristem cells, at least three randomly coded slides were observed under the light microscope. Calculation of the mitotic index was done by counting the number of dividing cells per total cells scored for each concentration. Different cell phase frequencies, mitotic index, and c-metaphase cells were analyzed by 2X2 contingency χ2-test. Pearson correlation was analyzed using GraphPad Prism 8.4.3.

## Results and discussion

The focus of the present investigation was to analyze cell cycle delay, metaphase arrest, and c-metaphase inducing effects of AQPEF in *A. cepa* root apical meristem cells. Many authors suggested that antiproliferative and cytotoxic effects of plant extracts or chemical substances can be evaluated using *A. cepa* root tip cells (Levan 1938, Bakare *et al*. 2000, 2012, 2013, Frescura *et al*. 2012). The root growth retardation depends on the antiproliferative potentials of the treated substances. Data indicxate that AQPEF induced a concentration-dependent *A. cepa* root growth retardation effect. The highest root length retardation percentage (68.51±0.56%, *p*<0.001) in 150 μg mL^−1^ at 72 h of AQPEF treatment and IC_50_ value of the root growth retardation was 23.68±5.5 μg mL^−1^ at 48h treatment (Table 1). Recently, Barman *et al.* (2020) and Das *et al*. (2021) reported that plant extract induced root growth retardation and the levels of root growth retardation were increased with the increasing extract concentration and suppression of cell division in *A. cepa* root tips (Murthy *et al.* 2011, Ray *et al.* 2013). For the determination of antiproliferative and cytotoxic potentials of synthetic or biochemicals including plant extracts, the *A*. *cepa* test model is widely used (Frescura *et al*. 2013, Khanna *et al*. 2013).

**Table 1:** Root growth retardation effect of AQPEF in *A. cepa.*

| AQPEF<br>($\mu\text{g mL}^{-1}$) | Root length (cm)<br>(Root length retardation %) | | | Growth rate (cm/24 h)<br>Growth rate retardation (%) | |
| --- | --- | --- | --- | --- | --- |
|  | 24 h | 48 h | 72 h | 48 h | 72 h |
| 0 | 3.08±0.10 | 4.32±0.10 | 5.51±0.11 | 1.24±0.10 | 1.19±0.03 |
| 12.5 | 2.57±0.07<br>(16.51±0.39) | 3.37±0.04<sup>b</sup><br>(22.00±0.83) | 3.73±0.10<sup>a</sup><br>(32.92±1.01<sup>a</sup>) | 0.79±0.06<sup>b</sup><br>(35.62±2.62<sup>b</sup>) | 0.36±0.11<sup>a</sup><br>(69.87±5.05<sup>a</sup>) |
| 25 | 2.38±0.08<sup>b</sup><br>(22.68±0.87<sup>b</sup>) | 2.80±0.08<sup>a</sup><br>(35.08±1.06<sup>a</sup>) | 3.09±0.12<sup>a</sup><br>(43.92±1.32<sup>a</sup>) | 0.42±0.04<sup>a</sup><br>(65.48±3.84<sup>a</sup>) | 0.28±0.06<sup>a</sup><br>(75.98±2.95<sup>a</sup>) |
| 50 | 2.17±0.07<sup>a</sup><br>(29.59±1.52<sup>a</sup>) | 2.45±0.09<sup>a</sup><br>(43.31±1.97<sup>a</sup>) | 2.59±0.18<sup>a</sup><br>(52.94±1.87<sup>a</sup>) | 0.28±0.16<sup>a</sup><br>(76.55±8.87<sup>a</sup>) | 0.14±0.03<sup>a</sup><br>(87.91±1.58<sup>a</sup>) |
| 100 | 1.80±0.08<sup>a</sup><br>(41.64±1.33<sup>a</sup>) | 1.99±0.07<sup>a</sup><br>(53.91±0.74<sup>a</sup>) | 2.13±0.07<sup>a</sup><br>(61.38±0.78<sup>a</sup>) | 0.19±0.03<sup>a</sup><br>(84.17±2.43<sup>a</sup>) | 0.13±0.02<sup>a</sup><br>(88.50±1.05<sup>a</sup>) |
| 150 | 1.50±0.06<sup>a</sup><br>(51.33±1.31<sup>a</sup>) | 1.63±0.05<sup>a</sup><br>(62.31±0.44<sup>a</sup>) | 1.73±0.05<sup>a</sup><br>(68.51±0.56<sup>a</sup>) | 0.13±0.03<sup>a</sup><br>(89.32±1.93<sup>a</sup>) | 0.10±0.03<sup>a</sup><br>(91.01±1.51<sup>a</sup>) |

*Allium cepa* root growth retardation effect of the AQPEF was performed in triplicate and data analyzed with Student's t-test. <sup>a</sup> significant at $p<0.001$ , <sup>b</sup> significant at $p<0.01$ and <sup>c</sup> significant at $p<0.05$ as compared to their respective control by Student's t-test. Data represented as Mean ± SEM.

The mitotic index percentage in root apical meristem cells varies with the growing periods and the used AQPEF concentrations. Here, the highest MI (13.25±0.72 %) was scored at 4 h of AQPEF (150 μg mL^−1^) treatment and the lowest MI (3.55±1.10 %) was found in the case of colchicine (150 μg mL^−1^) at 4 h followed by 16 h recovery samples. At 2 h, MI% values were gradually increased-5.67±0.30, 6.63±0.41, 7.58±0.5, 9.76±0.26, and 10.51±0.80 % respectively for 0, 50,100, 150, and 200 μg mL^−1^. The MI% also increased in the case of 4 h treated samples - 7.09±0.62, 10.3±0.56, 10.3±0.29, 13.25±0.72, and 9.95±0.80% respectively for 0, 50,100, 150, and 200 μg mL^−1^ of AQPEF. In the case of 16 h recovery samples (4 h T + 16 h RS), MI% values were 10.06±0.92, 7.00±0.29, 8.45±1.64, 6.51±0.77, and 7.43±0.38% respectively for 0, 50, 100,150 and 200 μg mL^−1^ of AQPEF. In the case of 150 μg mL^−1^ colchicine treatment, data indicate the MI% decreasing tendencies at 2 h (12.55±1.44%), 4 h (9.17±0.56%) and 4 h followed by 16 h recovery samples (3.55±1.10%). Plant extracts having mitostatic effects can be regarded as cytogenotoxic substance (Frescura *et al*. 2013, Khanna *et al*. 2013) and the reduction in the mitotic index is considered as an important parameter to examine the antimitotic as well as cytotoxic effects of biochemicals. The reduction in MI may be the result of their cytotoxic effects like blockage during DNA synthesis or G_2_ phase or insufficient synthesis of ATP during spindle fiber formation and elongation (Sreeranjini and Siril 2011).

The investigation by squash preparation of *A. cepa* root apical meristem cells revealed that AQPEF induced increased metaphase frequencies and decreased frequencies of prophase, anaphase, and telophase in a concentration-dependent manner at both 2 and 4 h treated samples indicate that AQPEF has definite metaphase arresting activity. Subsequently, in recovery root tip cells (4 h T + 16 h RS), the frequencies of prophase, anaphase, and telophase showed an increasing tendency as compared to 2 and 4 h treated cells (Table 2). The highest metaphase cells percentage (81.44±1.61%) was scored from 4 h AQPEF treated (100 μg mL^−1^) samples and that was followed by the concentration of 150 μg mL^−1^ (78.48±2.41%). However, in the case of 16 h recovery samples, the metaphase frequency showed a reducing tendency, and the metaphase frequencies were scored as 65.69±3.87 % and 52.42±6.02 % respectively for 100 and 150 μg mL^−1^ of AQPEF. Similarly. Colchicine treatment for 2 and 4 h notably induced increased metaphase frequency, 79.75±4.46%, and 84.88±1.26 % respectively. In the case of 16 h recovery after 4 h colchicine treatment, the metaphase frequency (38.81±1.66 %) was comparable to untreated control (Table 2). The mitotic index elevation in 2 and 4 h treated cells may be due to an increase in metaphase frequencies. A similar colchicine-induced change in mitotic index and metaphase frequencies was observed earlier (Davidson *et al*. 1966).

**Table 2:**
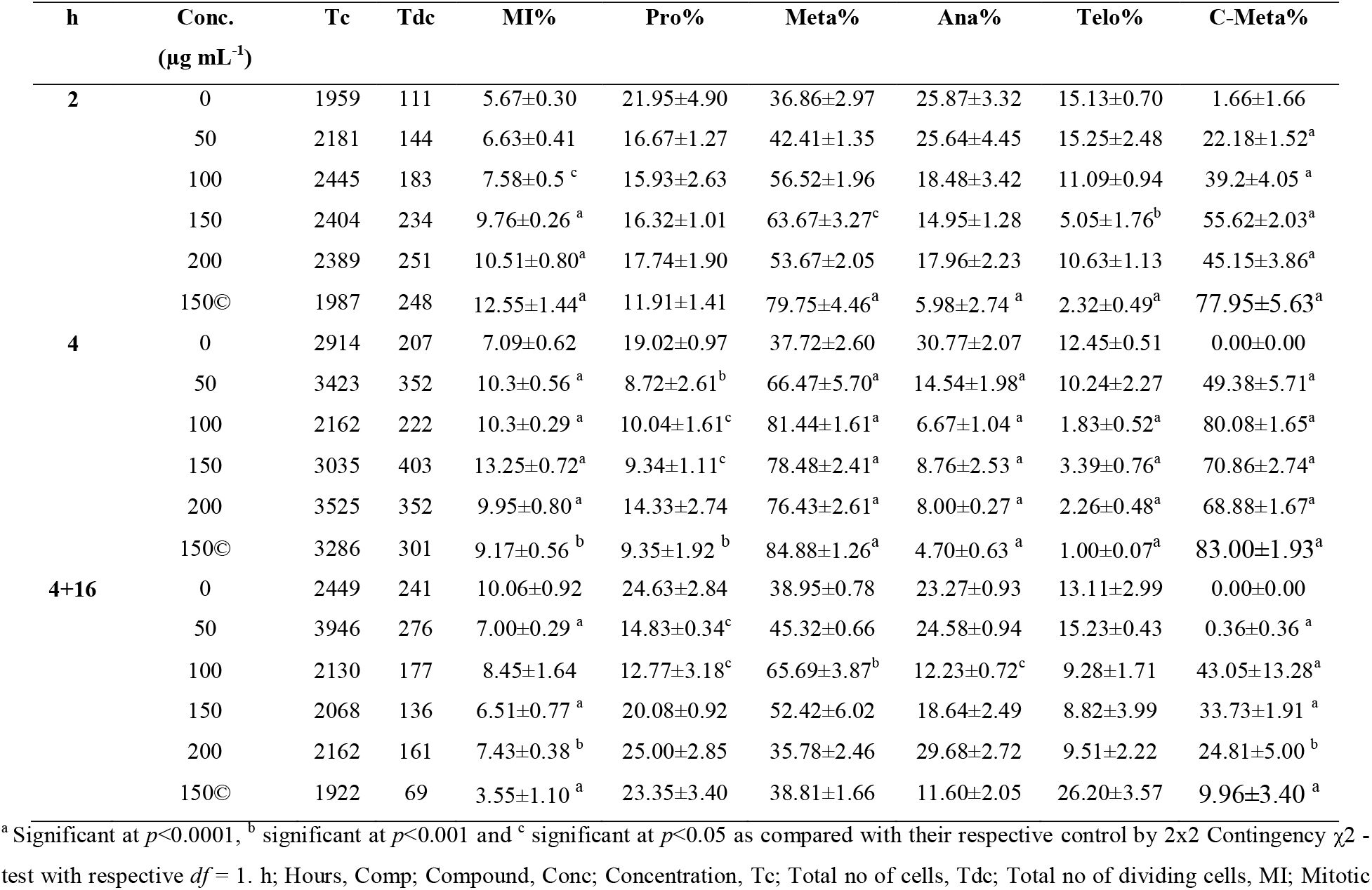
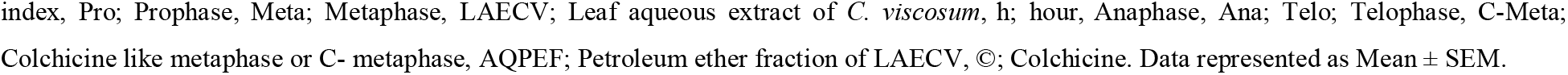
Metaphase arrest and c-metaphase inducing effects of AQPEF.

| h | Conc.<br>($\mu\text{g mL}^{-1}$) | Tc | Tdc | MI% | Pro% | Meta% | Ana% | Telo% | C-Meta% |
| --- | --- | --- | --- | --- | --- | --- | --- | --- | --- |
| 2 | 0 | 1959 | 111 | 5.67±0.30 | 21.95±4.90 | 36.86±2.97 | 25.87±3.32 | 15.13±0.70 | 1.66±1.66 |
| | 50 | 2181 | 144 | 6.63±0.41 | 16.67±1.27 | 42.41±1.35 | 25.64±4.45 | 15.25±2.48 | 22.18±1.52<sup>a</sup> |
| | 100 | 2445 | 183 | 7.58±0.5<sup>c</sup> | 15.93±2.63 | 56.52±1.96 | 18.48±3.42 | 11.09±0.94 | 39.2±4.05<sup>a</sup> |
| | 150 | 2404 | 234 | 9.76±0.26<sup>a</sup> | 16.32±1.01 | 63.67±3.27<sup>c</sup> | 14.95±1.28 | 5.05±1.76<sup>b</sup> | 55.62±2.03<sup>a</sup> |
| | 200 | 2389 | 251 | 10.51±0.80<sup>a</sup> | 17.74±1.90 | 53.67±2.05 | 17.96±2.23 | 10.63±1.13 | 45.15±3.86<sup>a</sup> |
| | 150© | 1987 | 248 | 12.55±1.44<sup>a</sup> | 11.91±1.41 | 79.75±4.46<sup>a</sup> | 5.98±2.74<sup>a</sup> | 2.32±0.49<sup>a</sup> | 77.95±5.63<sup>a</sup> |
| 4 | 0 | 2914 | 207 | 7.09±0.62 | 19.02±0.97 | 37.72±2.60 | 30.77±2.07 | 12.45±0.51 | 0.00±0.00 |
| | 50 | 3423 | 352 | 10.3±0.56<sup>a</sup> | 8.72±2.61<sup>b</sup> | 66.47±5.70<sup>a</sup> | 14.54±1.98<sup>a</sup> | 10.24±2.27 | 49.38±5.71<sup>a</sup> |
| | 100 | 2162 | 222 | 10.3±0.29<sup>a</sup> | 10.04±1.61<sup>c</sup> | 81.44±1.61<sup>a</sup> | 6.67±1.04<sup>a</sup> | 1.83±0.52<sup>a</sup> | 80.08±1.65<sup>a</sup> |
| | 150 | 3035 | 403 | 13.25±0.72<sup>a</sup> | 9.34±1.11<sup>c</sup> | 78.48±2.41<sup>a</sup> | 8.76±2.53<sup>a</sup> | 3.39±0.76<sup>a</sup> | 70.86±2.74<sup>a</sup> |
| | 200 | 3525 | 352 | 9.95±0.80<sup>a</sup> | 14.33±2.74 | 76.43±2.61<sup>a</sup> | 8.00±0.27<sup>a</sup> | 2.26±0.48<sup>a</sup> | 68.88±1.67<sup>a</sup> |
| | 150© | 3286 | 301 | 9.17±0.56<sup>b</sup> | 9.35±1.92<sup>b</sup> | 84.88±1.26<sup>a</sup> | 4.70±0.63<sup>a</sup> | 1.00±0.07<sup>a</sup> | 83.00±1.93<sup>a</sup> |
| 4+16 | 0 | 2449 | 241 | 10.06±0.92 | 24.63±2.84 | 38.95±0.78 | 23.27±0.93 | 13.11±2.99 | 0.00±0.00 |
| | 50 | 3946 | 276 | 7.00±0.29<sup>a</sup> | 14.83±0.34<sup>c</sup> | 45.32±0.66 | 24.58±0.94 | 15.23±0.43 | 0.36±0.36<sup>a</sup> |
| | 100 | 2130 | 177 | 8.45±1.64 | 12.77±3.18<sup>c</sup> | 65.69±3.87<sup>b</sup> | 12.23±0.72<sup>c</sup> | 9.28±1.71 | 43.05±13.28<sup>a</sup> |
| | 150 | 2068 | 136 | 6.51±0.77<sup>a</sup> | 20.08±0.92 | 52.42±6.02 | 18.64±2.49 | 8.82±3.99 | 33.73±1.91<sup>a</sup> |
| | 200 | 2162 | 161 | 7.43±0.38<sup>b</sup> | 25.00±2.85 | 35.78±2.46 | 29.68±2.72 | 9.51±2.22 | 24.81±5.00<sup>b</sup> |
| | 150© | 1922 | 69 | 3.55±1.10<sup>a</sup> | 23.35±3.40 | 38.81±1.66 | 11.60±2.05 | 26.20±3.57 | 9.96±3.40<sup>a</sup> |

<sup>a</sup> Significant at $p<0.0001$ , <sup>b</sup> significant at $p<0.001$ and <sup>c</sup> significant at $p<0.05$ as compared with their respective control by 2x2 Contingency $\chi^2$ - test with respective $df = 1$ . h; Hours, Comp; Compound, Conc; Concentration, Tc; Total no of cells, Tdc; Total no of dividing cells, MI; Mitotic
index, Pro; Prophase, Meta; Metaphase, LAECV; Leaf aqueous extract of *C. viscosum*, h; hour, Anaphase, Ana; Telo; Telophase, C-Meta; Colchicine like metaphase or C- metaphase, AQPEF; Petroleum ether fraction of LAECV, ©; Colchicine. Data represented as Mean ± SEM.

Data indicate that the C-metaphase is the most frequent type of abnormalities induced by AQPEF in *A. cepa* root apical meristem cells. The AQPEF and colchicine treatments for 2 and 4 h induced a significant (*p*<0.001& 0.0001) increase in the frequency of C-metaphase in *A. cepa* root apical meristem cells in a concentration-dependent manner and decreased in recovery treatments (Table 2, Fig. 1, 2). In the case of 2 h treatment, 150 μg mL^−1^ of AQPEF showed the highest C-metaphase frequency (55.62%) but in the case of 4 h treatment, 100 μg mL^−1^ of AQPEF induced the highest (80.08 %) C-metaphase. Similarly, C-metaphase frequency was also increased in colchicine (150 μg mL^−1^) treated samples as 77.95±5.63% and 83.00±1.93% respectively for 2 and 4 h. In the case of 16 h recovery samples (4 h T + 16 h RS), c-metaphase frequency (9.96±3.40%) was decreased. The formation of C-metaphase is directly correlated with the microtubule disruption and indicates that phytochemicals present in AQPEF may have colchicine-like microtubule destabilizing activity (FiskesjÖ 1985, Bonciu *et al*. 2018, Shahin and El-Amoodi 1991). These data also correlate with the occurrence of C-metaphase in LAECV and colchicine-treated *A. cepa* root tip cells (Kundu and Ray 2016). Recently, Barman *et al*. (2021) reported colchicine like *C. inerme* leaf aqueous extract, LAECI, induced haphazardly arranged condensed chromosomes, C-metaphase, in *A. cepa.* Correlation between the formation of C-metaphase, vagrant chromosome, and polyploidy was evident by many investigators in *A. cepa* root tip cells (Carvalho *et al*. 2019). Based on this correlative evidence, we can postulate that AQPEF has a similar effect to LAECV on *A. cepa* root tip cells.

**Fig. 1:**
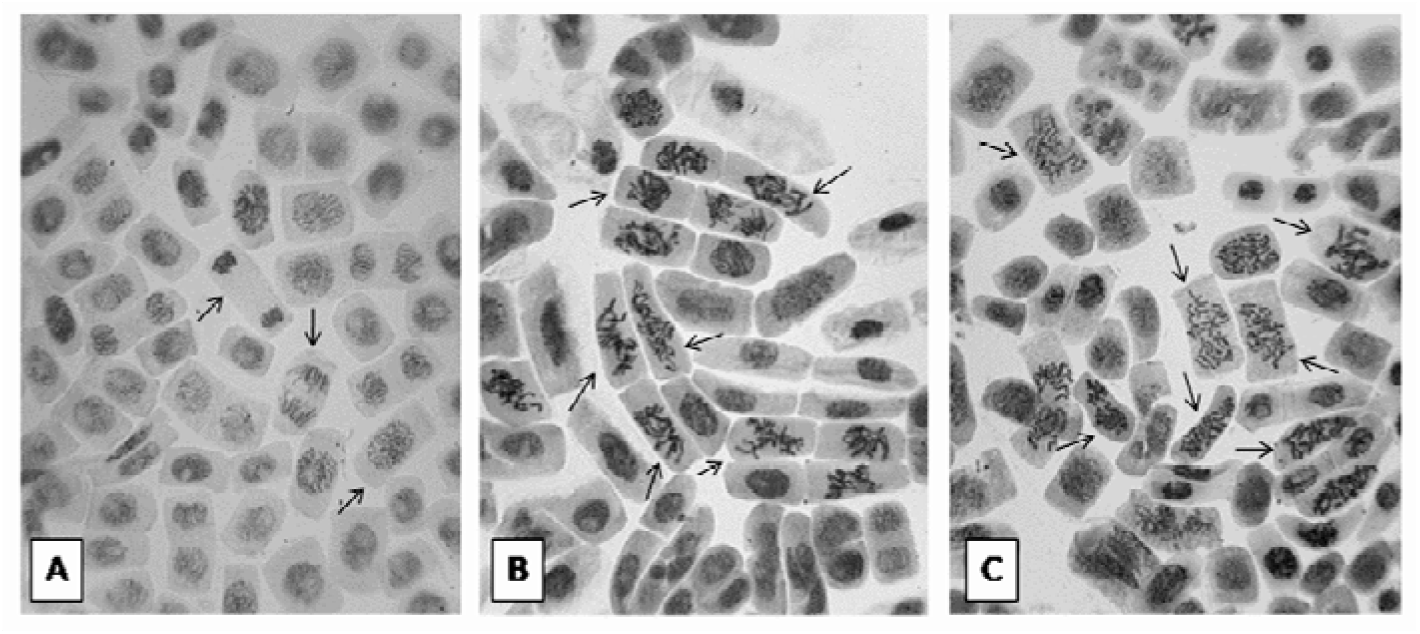
Photomicrograph showing the squash preparation of A, Untreated; B, AQPEF (petroleum ether fraction); C, Colchicine induced increased C-metaphase frequency in root apical cells of *A. cepa* at 4 h continuous treatment.

**Fig. 2:**
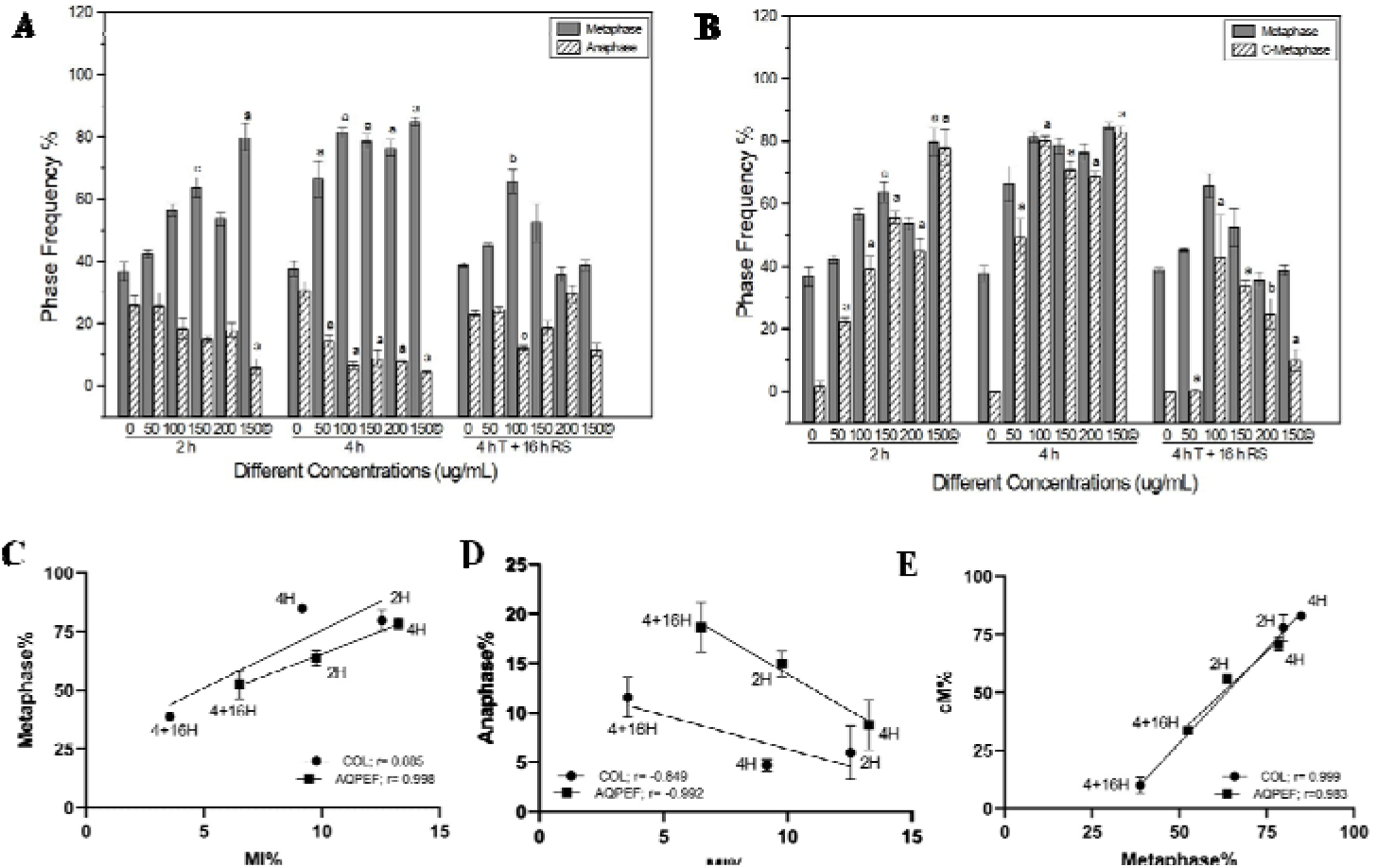
Graph showing the effect of AQPEF and Colchicine on A, metaphase and anaphase frequency; B, metaphase and C-metaphase frequency. Data represented as mean±sem. ^a^ significant at *p*<0.0001 and ^b^ significant at *p*<0.001, ^c^ significant at *p*<0.05 as compared to their respective control by 2×2 Contingency χ2 -test with respective d*f* = 1. Pearson Correlation and simple linear regression analysis of Colchicine (150μg mL^−1^) and AQPEF (150μg mL^−1^) between MI% (Mitotic Index%), META% (Metaphase%), ANA% (Anaphase%), and cM% (C-metaphase%) considering the change in their relative percentages at 2 h, 4 h and 4 h T+16 h RS. C, MI% and Metaphase%; D, MI% and Anaphase%; E, Metaphase% and cM%.

From Pearson correlation and simple linear regression analysis of colchicine (150 μg mL^−1^) and AQPEF (150 μg mL^−1^) between MI% (Mitotic Index%), META% (Metaphase%), ANA% (Anaphase%), and cM% (C-metaphase%) considering the change in their relative percentages at 2, 4, and 4h T+16h RS, a positive correlation was drawn between MI % and META %whereas a negative correlation was found between MI % and ANA % both in colchicine (r = 0.885 in META% and r = −0.849 in ANA%) and AQPEF (r = 0.998 in META% and r = −0.992 in ANA%). Positive correlation was also drawn between META% and cM% in AQPEF (r = 0.983) and colchicine (0.999). The above observations indicate that an increase in metaphase frequency was correlated with the decrease in anaphase frequency of both colchicine and AQPEF treatment. It can be extrapolated as the metaphase arresting nature of colchicine and AQPEF, leads to an increase in metaphase frequency and a simultaneous decrease in anaphase frequency, a possible mechanism for the cell cycle delay. A positive correlation between META% and cM% indicate that both compounds possess metaphase arresting activity; therefore, AQPEF may have colchicine-like microtubule destabilizing attributes (Fig.2). The LAECV and its petroleum ether fraction, AQPEF, has colchicine-like metaphase arrest and mitotic abnormalities inducing potentials (Ray *et al*. 2012, Kundu and Ray 2016, Roy *et al*. 2021) and this study indicates AQPEF has cell cycle delay, metaphase arrest, c-metaphase inducing potentials in *A. cepa* root apical meristem cells. A further detailed investigation is required for purification of active principles of *C. viscosum* and to test their relative cell cycle delay, pro-metaphase arrest, and mitotic abnormality-inducing potentials.

## Acknowledgments

The authors acknowledge Prof. A. Mukherjee for plant species authentication and the financial support of UGC-SRF (FC(Sc)/RS/UGC/ZOO/2018-19/129, w.e.f. 07.04.2018, dated: 04.02.2019), and the DST-PURSE, DST-FIST, and UGC-DRS-MRP-sponsored facilities in the Department of Zoology.

